# Increased copy number couples the evolution of plasmid horizontal transmission and antibiotic resistance

**DOI:** 10.1101/2020.08.12.248336

**Authors:** Tatiana Dimitriu, Andrew Matthews, Angus Buckling

## Abstract

Antimicrobial resistance (AMR) in bacteria is commonly encoded on conjugative plasmids, mobile elements which can spread horizontally between hosts. Conjugative transfer disseminates AMR in communities but it remains unclear when and how high transfer rates evolve, and with which consequences. Here we studied experimentally the evolution of two antibiotic resistance encoding plasmids when confronted to different immigration rates of susceptible, plasmid-free hosts. While plasmid RP4 did not evolve detectably, plasmid R1 rapidly evolved up to 1000-fold increased transfer rates in the presence of susceptible hosts, at a cost to its host. Unexpectedly, most evolved plasmids also conferred to their hosts the ability to grow at high concentrations of antibiotics. The most common mutations in evolved plasmids were contained within the *copA* gene which controls plasmid replication and copy number. Evolved *copA* variants had elevated copy number, leading to both higher transfer rates and AMR. Due to these pleiotropic effects, host availability and antibiotics were each sufficient to select for highly transmissible plasmids conferring high levels of antibiotic resistance.

## Introduction

Conjugative plasmids are mobile genetic elements that transmit horizontally within and between species of bacteria ^1^. Plasmids often confer antimicrobial resistance (AMR) to their hosts, hence understanding the ecological and genetic determinants of variation in transmission rate (transfer) among plasmids ^2^ and host bacteria ^3^ is central to managing the evolution of AMR ^4–6^. While greater transfer is expected to increase the frequency of resistant bacteria in a population, there may be less predictable correlated effects on AMR. For example, plasmids that are at high frequencies within communities may rapidly adapt to ameliorate the potential costs ^7^ they impose on their hosts in the absence of antibiotic selection, further increasing their frequency ^8–10^. Transfer rates might also correlate with the level of AMR conferred to the host cell, for instance if transfer trade-offs with host growth ^11^, promotes loss of AMR genes ^12^, or is mechanistically linked to AMR ^13^. Here, we experimentally investigate the potential correlated consequences of selection for increased plasmid transfer rates on AMR.

Increased plasmid transfer (or generally parasite transmission) is predicted to only be favoured when susceptible hosts are present in abundance ^14^. This is because conjugative transfer imposes a cost to host bacteria. This cost in turn limits plasmid vertical transmission, leading to a trade-off between horizontal and vertical transmission ^11^. Indeed, evolution experiments in conditions with low opportunity for transfer observe decreased transfer rates and carriage costs ^9,15–17^, but horizontal transmission can increase when susceptible hosts are available ^11,18–20^. We evolved two conjugative, multidrug-resistant plasmids, R1 and RP4 that differ in their transfer regulation ^21,22^, under variable susceptible host availability, and followed the evolution of transfer rates and AMR. We conducted molecular analyses to understand the mechanisms underpinning our results, and hence their potential generality.

## Results

### R1 plasmid evolves high transfer rates in the presence of susceptible hosts

To vary the importance of horizontal transmission in plasmid life cycle without enforcing direct selection for plasmid-bearing cells, we passaged an initial culture of plasmid-bearing cells (either plasmid RP4 or R1) with regular influx of various proportions of immigrant plasmid-free cells (Figure 1A). We used both wild-type (wt) and mutator (mut) *Escherichia coli* host; the latter to increase genetic variation available for evolution. For both plasmids, maintenance in immigration treatments required horizontal transmission, or they would be rapidly diluted out, and plasmids in high immigration treatments experienced higher selection for horizontal transmission (Figure S1). RP4 plasmid was rapidly lost from all treatments with ≥ 90% immigration per day, but stably maintained over 30 days, after an initial decline, under 68% immigration. R1 plasmid was maintained for longer but R1-bearing cell density decreased steadily for all treatments, with a faster decrease under higher immigration (Figure 1B).

**Figure 1:**
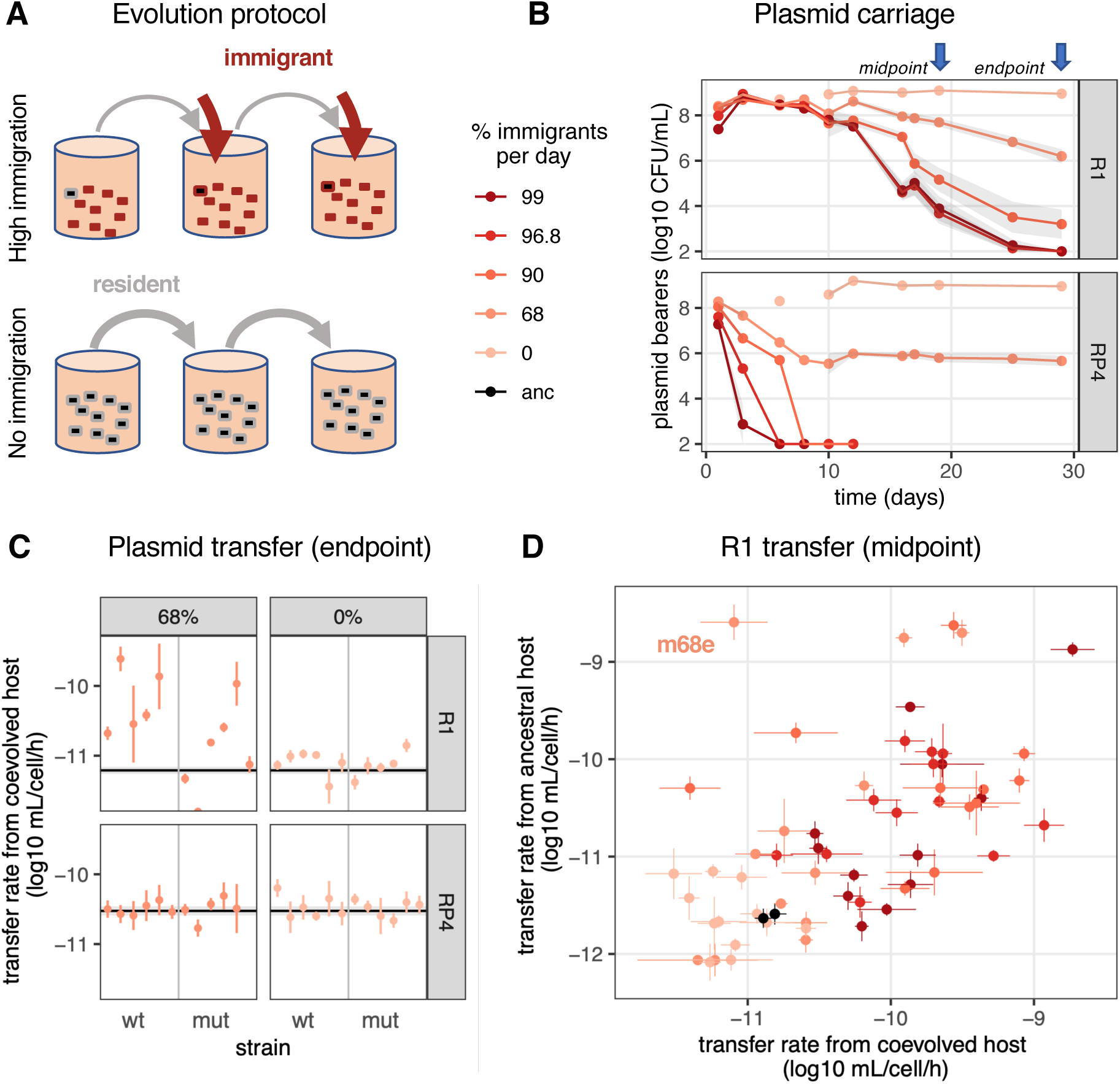
Evolution of plasmid transfer rate with varying host availability. A) Experimental evolution regime. Plasmid-bearing cells (black) were first mixed with plasmid-free cells, then evolving populations were diluted every day to fresh medium (gray arrows) mixed in varying frequencies to plasmid-free cells (red arrows). Each treatment was performed in 6 replicates with either wt or mut *E. coli* hosts. B) Plasmid population dynamics. Plasmid carriage was estimated by measuring ampicillin resistance. Initially one of six replicates was measured until day 12; then the treatment average is shown ± s.e.m. (shaded area). C) Endpoint clone conjugation rates. The black line and shaded area are respectively the geometric mean and standard error (SE) of ancestral plasmid transfer rate; each coloured dot and line indicates respectively the geometric mean and SE of evolved clones (N=3). D) Midpoint R1 conjugation rates. The x axis shows transfer rates measured from plasmid-bearing coevolved hosts; the y axis from ancestral hosts carrying evolved plasmids. Each coloured dot and line indicate respectively the geometric mean and SE of evolved clones (N=3); black dots and lines show ancestral plasmids.

We first measured transfer rates from coevolved plasmid-host combinations to ancestral recipients. Using one plasmid-bearing clone per lineage as a donor and a standard plasmid-free recipient, we detected no significant changes in endpoint (29 days) conjugation rates for RP4, but increased transfer for R1 with 68% immigration (Figure 1C, R1 transfer rate ∼ immigration, F_2,86_=5.3, *p*=0.007). For R1, we next focused on midpoint clones, before plasmid lineages under high immigration went extinct. We observed significant effects of immigration treatment on evolved transfer rates (Figure 1D, transfer rate ∼ immigration, F_5,198_=42.6, *p*<2.10^−16^). Specifically, treatments with ≥90% immigration had significantly increased transfer rates compared to the ancestor, no immigration and 68% immigration treatments (Tukey test, *p* < 2.10^−5^) but were not significantly different from each other. Thus, high transfer rates evolved in the presence of abundant (≥90% immigration) plasmid-free recipients.

The above assay determined the net effects of host and plasmid (co)evolution in transfer. To tease out host effects, we also measured transfer rates of midpoint evolved plasmids from the ancestral host. Transfer rates were still increased in plasmids from immigration treatments (Figure S2A). Transfer rates from coevolved hosts correlated with the ones from unevolved hosts (Figure 1D, bivariate model Cov(coevolved, unevolved host)=0.39±0.10, clone covariance effect *χ*=23, *p*=1.5.10^−6^; correlation coefficient=0.60±0.09). Thus, increased conjugation rates are mostly due to genetic changes on the plasmid. However, a few clones diverged from this pattern (Figure 1D), most significantly clone mut68e, with a 1000-fold increase in transfer rate when the plasmid was present in an ancestral host, but not in its coevolved host. This suggests that some hosts evolved repression of plasmid transfer.

### R1 evolves higher cost but also increased AMR

Host repression of transfer suggests that transfer is deleterious to the host. As a proxy for plasmid cost, we measured the density donor hosts achieve during conjugation assays. Donor density was negatively correlated with transfer rates (Figure 2A, donor density ∼ transfer rate, estimate=-0.52±0.03, r^2^=0.55, *p*<2.2 10^−16^). Host fitness still correlated with transfer rates when plasmids were carried by the same, ancestral host (Figure S2B), confirming that some of the reduction in cell growth is due to plasmid evolution. This trade-off will limit the fitness benefit of increased transfer rate, by limiting plasmid vertical transmission. Accordingly, high plasmid densities during evolution were not correlated to high clone transfer rate (Figure S3).

**Figure 2:**
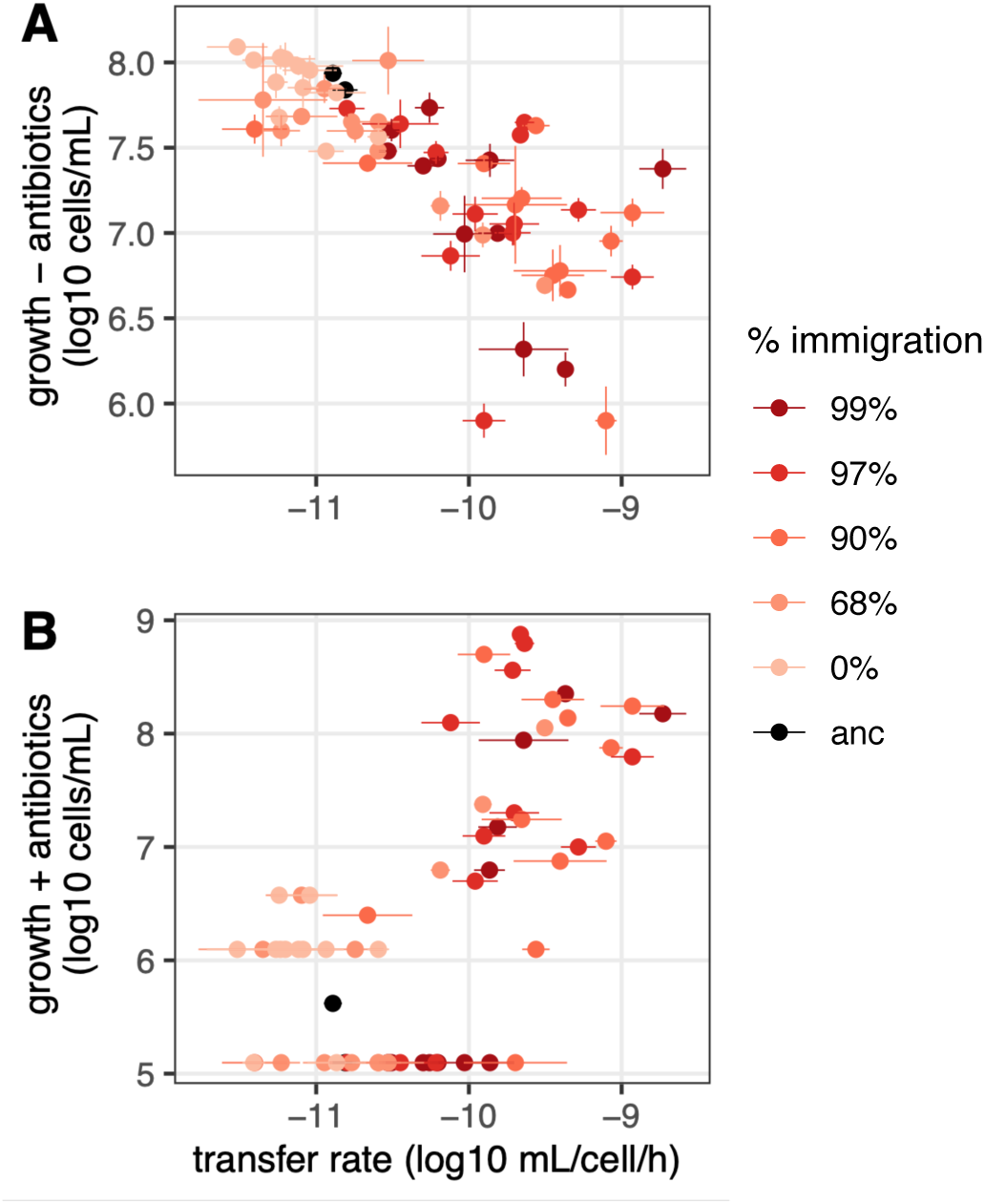
Breakdown of transfer-host fitness trade-off under antibiotic selection. Host growth in the absence of antibiotics (A, donor cell density during conjugation assays) and in the presence of high levels of antibiotics (B, density of cells able to grow on Amp 0.5g/L) is shown for midpoint coevolved clones as a function of their transfer rate. Dots and lines indicate respectively the geometric average and standard error (N=3).

To evaluate if the benefits conferred by plasmids in the presence of antibiotics also evolved, we measured cell growth in the presence of high levels of antibiotics (Figure 2B). Many evolved clones conferred to their host an ability to grow in conditions in which the ancestor could not (Figure 2B). Most of these also had increased transfer rate (Spearman rank-correlation *ρ =* 0.62, *p* = 10^−7^), suggesting that increased AMR is a side-effect of selection for transfer.

This apparent coupling between high transmission and AMR prompted us to ask if selection for one trait can promote the other. First, we plated early populations from the evolution experiment (9 days) directly on Amp 0.5g/L. Highly resistant cells were more common in immigration treatments, especially in the mutator host (mut) (log_10_ (cell density) ∼immigration * host, F_1,56_=12.8, *p*=0.0007), (Figure 3A). In some lineages, nearly all plasmids conferred upon their hosts ability to grow at high antibiotic concentrations, and almost all showed increased transfer compared to R1_wt_ (Figure S4). Thus, immigration treatments selecting for increased transfer collaterally promote the appearance of high AMR.

**Figure 3:**
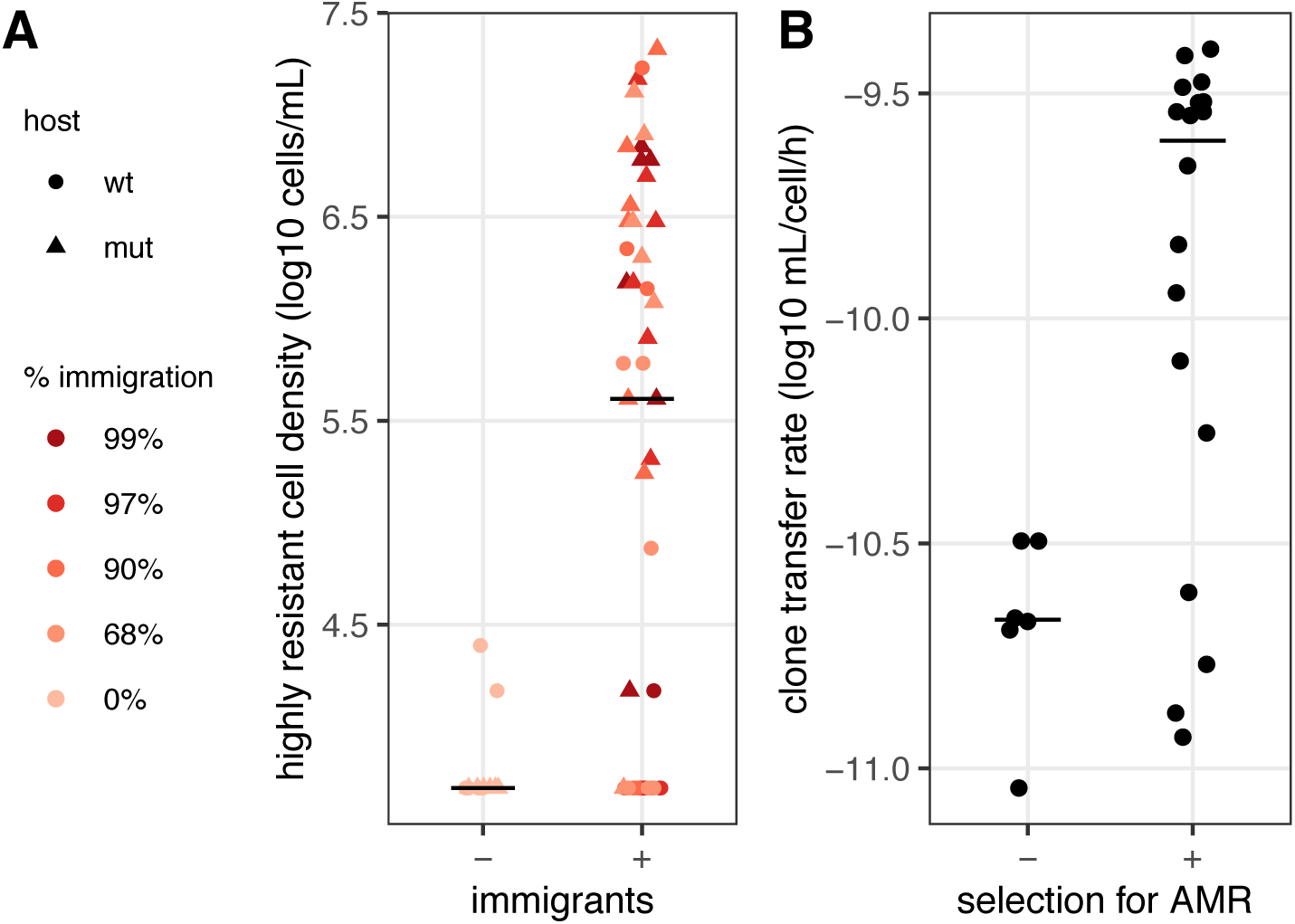
Reciprocal selection for high transfer rate and AMR. A: Host availability promotes the evolution of highly antibiotic-resistant plasmids. The density of cells able to grow in the presence of 0.5g/L Amp is shown for populations evolved for 9 days with or without immigration. B: Selecting for high level antibiotic resistance promotes evolution of highly transferable plasmids. 6 independent populations of a standard host carrying R1_wt_ were plated with or without high concentrations of antibiotics, and the transfer rate of 3 spontaneous mutants per population measured (geometric average of N=3 replicates per mutant). Horizontal lines show the median value for each treatment.

Conversely, selecting for high levels of resistance might also increase conjugative transfer rates in the absence of direct selection for transmission. We plated independent cultures of the ancestral host carrying R1_wt_ on high levels of antibiotics, and measured plasmid transfer rate for highly resistant spontaneous mutants (Figure 3B). Selection for AMR increased transfer rate overall (log_10_ transfer rate ∼ selection, F_1,70_=30.6, *p*=5.10^−7^), and 14 of 18 mutants displayed increased transfer compared to R1_wt_, with approximately 10-fold higher transfer rates (Figure 3B).

### Increased transfer and AMR are coupled via an increase in R1 copy number

To understand the genetic basis of phenotypic changes and their coupling, we sequenced the previously characterised midpoint evolved clones. A summary of mutations is shown in Table S2. Plasmid mutations were found almost exclusively in clones from immigration treatments (Figure 4A). Four evolved plasmids from the 90% immigration treatment contain large deletions (∼10 to 20 kb), which include one or several antibiotic resistance determinants (Figure S5). Next, several plasmid loci present parallel mutations in independently evolved lineages, a sign of convergent molecular evolution. Three evolved plasmids carried the same frameshift in the coding region of *finO*, the main repressor of the transfer operon ^21^, and another carried a 7bp insertion. These mutations are known to cause loss of function and derepression of the transfer operon ^23^. 16 variants had a G deletion in a polyG tract ∼ 50bp before the coding region of *gene32*, part of the leading region first transferred to recipient cells. *gene32* function is unknown but it is conserved and likely important for conjugative transfer ^24^. Finally, the most common mutations were in the *copA* locus. The untranslated antisense CopA RNA plays a central role in R1 replication and copy number control by inhibiting translation of the initiator for plasmid replication RepA ^25^. We observed 27 point mutations variants contained in a 13bp region within *copA*, all from C or G nucleotides towards A or T, denoted as *copA\** below (Figure 4A). These mutations structurally alter the head of CopA RNA’s loop II that is involved in binding to RepA mRNA ^25^. Identical or similar mutations were shown to increase R1 plasmid copy number (PCN) ^25^ by decreasing CopA binding to RepA mRNA ^26^. Accordingly, we observed increased sequencing coverage for *copA\** plasmids consistent with increased PCN (Figure 4B top, detail in Figure S5).

**Figure 4:**
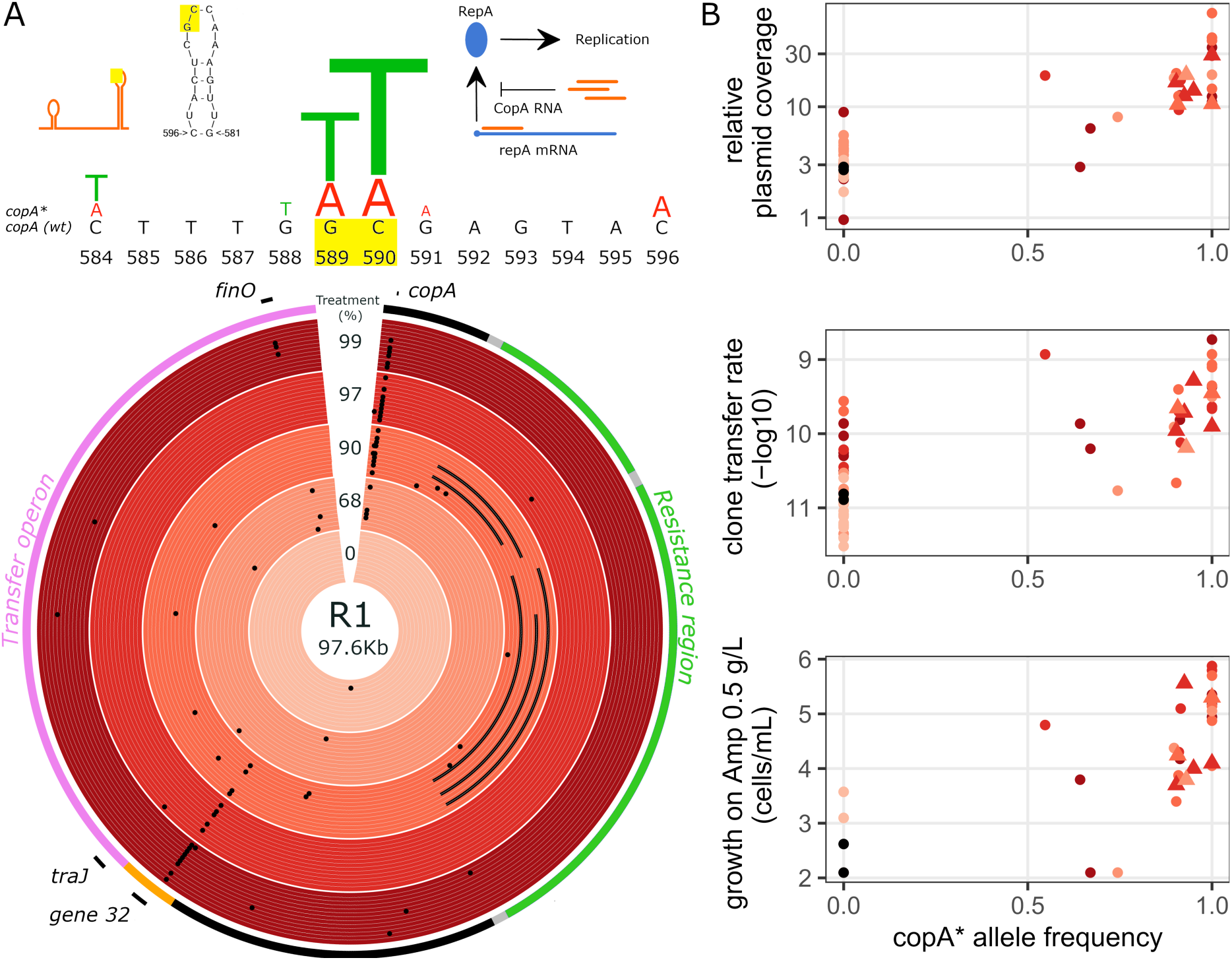
Molecular evolution and mechanistic basis for increased R1 transfer and AMR. A: Variants detected in evolved plasmids. The outermost ring of the circos plot shows a simplified plasmid map. Evolved plasmids are then shown by treatment, from outside in wt a-f then mut a-f clones. Black dots indicate small polymorphisms, lines show large deletions. Inset diagrams above show the secondary structure of CopA antisense RNA and its mode of action. Significant 13bp of evolved variants are summarised with letter size proportional to the relative abundance of each mutation. The position of the most common mutations in the second loop of CopA RNA is highlighted in yellow (modified from ^25^). B: Phenotypic effects of *copA\** variants. Evolved plasmid coverage, transfer rate and growth on Amp 0.5g/L are shown as a function of *copA\** mutation frequency in the evolved clone (as not all plasmid copies necessarily have *copA\** mutation). Triangles indicate clones for which no other mutation than *copA\** was present on the plasmid.

Across all clones, only *copA\** and *gene32* mutations were significantly associated to high transfer rates, *copA\** having the strongest effect (log_10_ transfer rate ∼ *copA\**+ *gene32* + *finO, copA\** effect 1.02±0.15, F_1,200_=235, *p*<2.10^−16^, *gene32* effect 0.30±0.17, F_1,200_=11.8, *p*=0.0007). Moreover, seven evolved plasmids carried only a *copA\** mutation and were otherwise identical to R1_wt_ (triangles in Figure 4B); these had on average a 17-fold increased transfer rate (*copA\** effect, F_1,97_=171, *p*<2.10^−16^) and a 55-fold increase in AMR (F_1,25_=45, *p*=5.10^−7^). These plasmids also had 18-fold increased transfer rate when in the ancestral host (Figure S6, F_1,109_=160, *p*<2.10^−16^). Thus, *copA\** mutations are sufficient to confer a 17-fold increase in transfer rate, and explain (via increased PCN) its coupling with AMR.

High-level AMR linked to increased PCN is explained by increased gene expression from plasmid-carried resistance genes ^27,28^. Correlated increase in transfer rate could happen in two ways: increased expression of the transfer operon (which encodes proteins responsible for transfer), or increased number of copies of the origin of transfer, *oriT* (on which the transfer machinery assembles to initiate transfer). An increase in *oriT* copy number will only increase transfer of the evolved R1 replicon itself, whereas increased protein expression should also increase mobilisation of other molecules carrying *oriT*. We found that a *finO* variant (with derepressed transfer expression ^23^) increases strongly transfer of both itself and pMOB, a non-conjugative plasmid carrying *oriT*, by similar ratios (Figure S7; R1_*finO*_ 608-fold, pMOB 648-fold, paired two-tailed t-test, *p*=0.85). A *copA\** variant also increased transfer for both plasmids but mobilised pMOB ∼4-fold less than itself (R1_*CopA\**_ 27-fold, pMOB 6.5-fold, *p*=0.0003). Thus, increases in transfer operon expression and in available *oriT* sequences both contribute to the increased transfer of *copA\** variants.

Finally, we compared in detail the effects of one representative *copA\** variant and a *finO* variant (Figure S8). In the absence of selective antibiotics, R1_*copA\**_ conferred a fitness cost to its host when competing with R1_wt_, but this cost was much lower than R1_*finO*_ cost (relative fitness W_*copA\**_=0.73±0.06, W_*finO*_=0.32±0.14, two sample t test t=-6, *p*=3.10^−5^). At high doses of antibiotics to which R1 confers resistance, R1_*copA\**_ increased cell survival compared to R1_wt_ and R1_*finO*_. In the presence of antibiotics to which plasmid-bearing cells are particularly sensitive ^27^, R1_*copA\**_ also reduced survival less than R1_*finO*_ (Figure S8).

## Discussion

In the presence of susceptible hosts, R1 plasmid rapidly evolves increased horizontal transmission. In most clones, increased transfer is linked to increased AMR due to mutations affecting the CopA RNA controlling PCN. Contrary to our expectations, few clones evolved derepression of the transfer operon, a previously well-characterized mechanism ^23^. This is likely due to the strong cost of full derepression. Indeed, high transfer rates traded off strongly with host growth across clones. We also observed some loss of resistance determinants, which might alleviate part of plasmid cost in the absence of antibiotics. Rapid loss of resistance genes by R1 was observed previously ^12,16^, and is likely underestimated here as we used Amp resistance as a proxy for plasmid carriage.

PCN could evolve by the same mechanism in other plasmids, as many plasmids regulate their PCN through antisense RNAs ^29^, and similarly point mutations are sufficient to change PCN ^28,30,31^. Increase in mobilisation when more *oriT* sequences are present is likely a general phenomenon for both conjugative and mobilisable plasmids. Increased transfer operon expression might be less widespread and depend on plasmid regulatory networks. For instance, in contrast to rapid R1 evolution, we observed no evolution in RP4 transfer rate: its regulation network, being based on repression rather than activation, might make transfer rate less evolvable ^21,22^.

In addition to increased horizontal transmission, *copA\** mutations also increased AMR through increased gene dosage. High levels of AMR correlated with high PCN have been observed previously ^25,27,28,31^; we show here this can be driven by selection for transfer. Interestingly, specific coupling between horizontal transmission and AMR has been observed in other mobile elements ^13,32^, due to mechanistic links between expression of transfer and resistance genes. The basis for coupling here is less specific, as all genes carried on the plasmid will experience higher dosage. Thus, other plasmid-encoded phenotypes might be affected similarly. For instance, increased PCN during infection controls virulence in *Yersinia*, via changes in CopA RNA levels ^33^.

Increased PCN could also make plasmid-encoded traits more evolvable. Multicopy plasmids have increased mutation supply, which can promote the evolution of AMR by point mutations ^31^. Indeed, *copA\** variants have significantly more additional SNPs than other evolved variants (Wilcoxon rank sum test, W=238, *p*=0.0009), suggesting that evolution of PCN is sufficient to rapidly increase evolvability. However, high PCN can also limit evolvability because it affects both genetic drift and selection acting on plasmid mutations ^34^. Selection will further depend on the genetic dominance of plasmid alleles ^35^. *copA\** variants might differ in all these aspects, as they will have high PCN but also experience frequent bottlenecks due to conjugation.

Classically, selecting for cells containing plasmids kills plasmid-free cells, removing plasmid-free hosts available for transfer and thus decreasing horizontal transmission overall ^36^. We show here that selection for high gene dosage of plasmid-carried genes can favour highly transmissible plasmids, because of the pleiotropic effects of increased PCN. This confirms that when horizontal transmission and plasmid benefits are linked, limiting opportunities for horizontal transmission can actually limit the evolution of mutualism ^37^. Other factors will likely shape this coupled evolution of PCN and horizontal transmission. On the plasmid side, PCN control is itself subject to conflicting selective pressures, with intracellular selection promoting selfishness, and intercellular selection promoting restraint ^38,39^. On the host side, repression of transfer and PCN alike will limit plasmid costs, but selection for plasmid genes might also favour alleles promoting transfer if plasmids are transferred to kin ^40^. The coupling of horizontal transmission with PCN and its pleiotropic effects might modify all these co-evolutionary outcomes, and consequently complicate AMR management.

## Methods

### Strains, plasmids and growth conditions

The ancestral plasmid-bearing strains were *E. coli* MG1655 (wt) and MG1655 *mutL*::KnR (mut) obtained by transduction from the Keio collection ^41^. Plasmids R1 and RP4 were conjugated into ancestral strains from the MFDpir donor strain ^3^. Plasmid-free immigrants were variants of wt and mut strains marked with *td-Cherry* ^40^ for transfers 1 to 15 and with a Δ*lac* deletion ^42^ for transfers 16 to 29. Cells were grown in LB medium. Plasmid-bearing clones were selected with Amp 100mg/L. Spontaneous rifampicin-resistant (Rif^R^) and nalidixic acid-resistant (Nal^R^) mutants of MG1655 ^3^ were used respectively as a standard donor host for measuring plasmid transfer from an ancestral genotype, and as a standard recipient in conjugation assays, using Rif 100 mg/L and Nal 30 mg/L for selection. Plasmids were transferred to plasmid-free backgrounds by conjugation, then transconjugants were selected on Amp + appropriate selection for the recipient phenotype. High-level evolved resistance to Amp was measured by plating on LB-agar + Amp 0.5g/L; when no colonies were detected a threshold value was calculated assuming one colony at the lowest dilution plated. Spontaneous mutants with increased antibiotic resistance using R1-carrying cells were obtained by plating 100 µL overnight cultures on Amp 500 mg/L + streptomycin (Str) 200 mg/L.

### Evolution experiment and evolved clones

Evolving populations were grown in 200 µL LB medium in 96-well plates, at 37C with 180rpm shaking and 100-fold overall dilution every 24h. For immigration treatments, plasmid-free immigrants were grown fresh from glycerol stock and mixed with the evolving resident cultures in the stated ratios at each passage. Plasmid-free *td-Cherry* strains were also passaged with 100-fold daily dilution. 6 replicate lineages were evolved per treatment x host strain (wt or mut) combination. One individual clone per lineage was picked for characterisation at various timepoints (see Table S1 for detail of midpoint clones).

### Conjugation assays

Plasmids-carrying donors and MG1655 Nal^R^ were grown overnight without antibiotic selection. Donor strains were first diluted 5-fold into LB medium and grown at 37C for 1h, then 20 µL diluted donor cultures were mixed with 20µL recipients and 160µL pre-warmed LB. After 1h mating, serial dilutions were plated on Amp, Nal and Amp+Nal to estimate densities of donors + transconjugants, recipients and transconjugants respectively. Conjugation rates and threshold values when no transconjugant was detected were calculated as described in ^3^. Replicates from the ancestral host assay (Figure 2A y-axis) with donor densities < 4.10^8^ cells/mL or no transconjugants were discarded as they were associated to evaporation during the conjugation assay.

### Sequencing and bioinformatics

Clones selected for sequencing (Table S1) were grown overnight, with Amp selection for plasmid-carrying clones, and DNA was extracted using CTAB extraction. Illumina whole-genome sequencing was provided by MicrobesNG (http://www.microbesng.uk). Sequencing data were mapped to a reference genome combining MG151655 (GenBank accession NC_000913) and R1 plasmid (GenBank accession KY749247), using breseq 0.35.1 ^43^, run in polymorphism mode to account for the coexistence of several plasmid alleles in high PCN clones. Variants not present at frequency > 20% in at least one clone, and variants also detected in ancestral clones were discarded. Large deletions were called manually using coverage data, generated with breseq command bam2cov with default settings. Circos plots were made with R package omiccircos ^44^.

### Mobilisation experiment

R1 *oriT* was amplified using primers 5’-CACGAAGCTTGCCTGCACTTTCGCCATATG-3’and5’-CACCGAATTCAATCAGTGGCCTGGCAGATC-3’, then cloned into pHERD30T plasmid ^45^, using restriction enzymes HindIII and EcoRI. The resulting plasmid pMOB was transformed into MG1655. pMOB-bearing cells were selected using 50 mg/L gentamycin (Gen), a dose necessary to avoid low-level resistance conferred by R1 variants. Clones containing both pMOB and a R1 variant were obtained by conjugation of R1 into MG1655 + pMOB using Amp+Gen selection. Mating was conducted as described above, but donor clone overnight cultures contained Amp+Gen, and mating time was reduced to 30min to limit secondary transfer; cells were also plated on Gen and Gen+Nal.

### Plasmid competition assays

After overnight growth without antibiotics, strains were mixed and grown for 24h after 100-fold overall dilution in 96-well plates. Densities were estimated immediately after dilution and after 24h, by plating serial dilutions on selective medium. Competition assays were run with equal ratios of plasmid-bearing cells, using two-way competitions with *td-Cherry* and Δ*lac* marked MG1655 hosts (N=6 for each combination, results with opposite host markers were pooled).

### Antibiotics effects

Plasmid effect on survival in the presence of antibiotics was measured with plasmids carried by MG1655Δlac. Each strain was grown overnight in 8 independent replicates, without antibiotics, then dilution series were plated on agar with various concentrations of antibiotics. Survival was defined as colony-forming units (CFUs) on antibiotic medium / CFUs on LB agar without antibiotics. To survey growth in the presence of antibiotics for all evolved midpoint clones, cultures of evolved were first grown overnight in LB + Amp 100mg/L, then plated on agar with Amp 0.5 g/L.

### Data analysis

Statistical analysis used R ^46^. Transfer rate and cell density values were log-transformed before analysis. For conjugation assays, data with donor densities < 4.10^5^cells/mL were excluded, as well as transconjugant densities = 0, which corresponded to replicates with high evaporation. To analyse the effect of host background on clone transfer rate (Figure 2A), where data from each background were obtained in separate experiments, a bivariate model was run using the ASReml package with clone as a random effect ^47^.

## Acknowledgments

We thank Alastair Wilson for help with statistical analysis.

## Notes

### Competing Interest Statement

The authors have declared no competing interest.

