## Supplemental figures for "Increased copy number couples the evolution of plasmid horizontal transmission and antibiotic resistance"

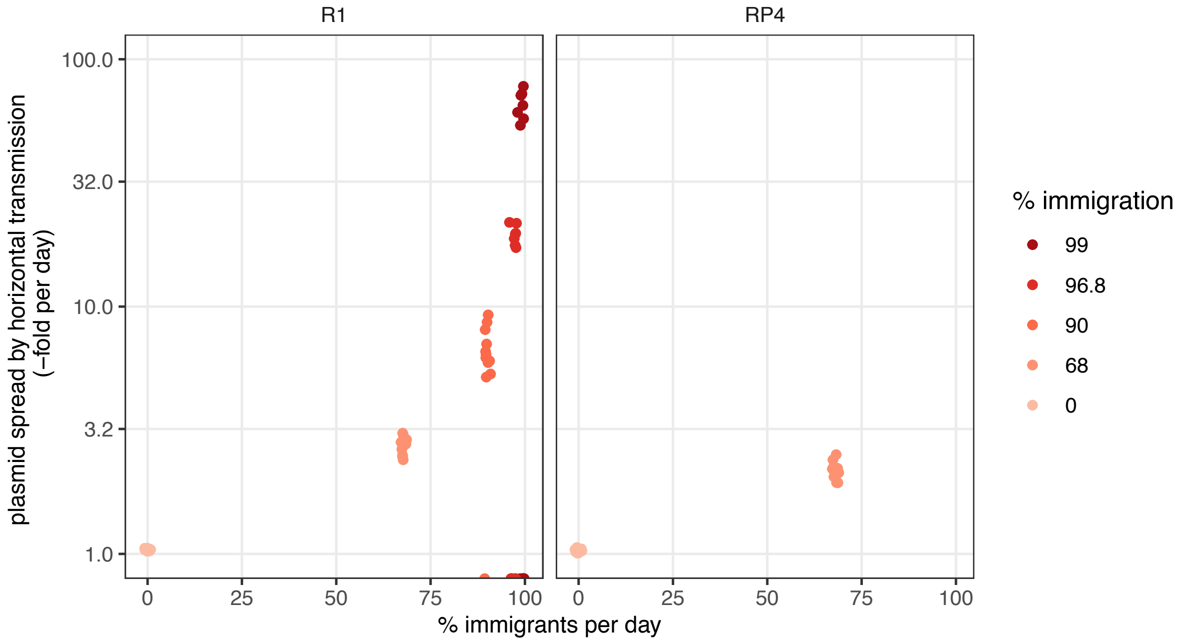


**Figure S1: Average plasmid amplification per day.** Theoretical plasmid-bearing cell densities with only vertical transmission were calculated after 19 days assuming equal fitness for all hosts. Plasmid spread by horizontal transmission was defined as the average daily increase in plasmid densities necessary to explain the deviation of measured densities at day 19 from theoretical ones in the absence of horizontal transmission. Zero values indicate lineages with plasmid extinction.

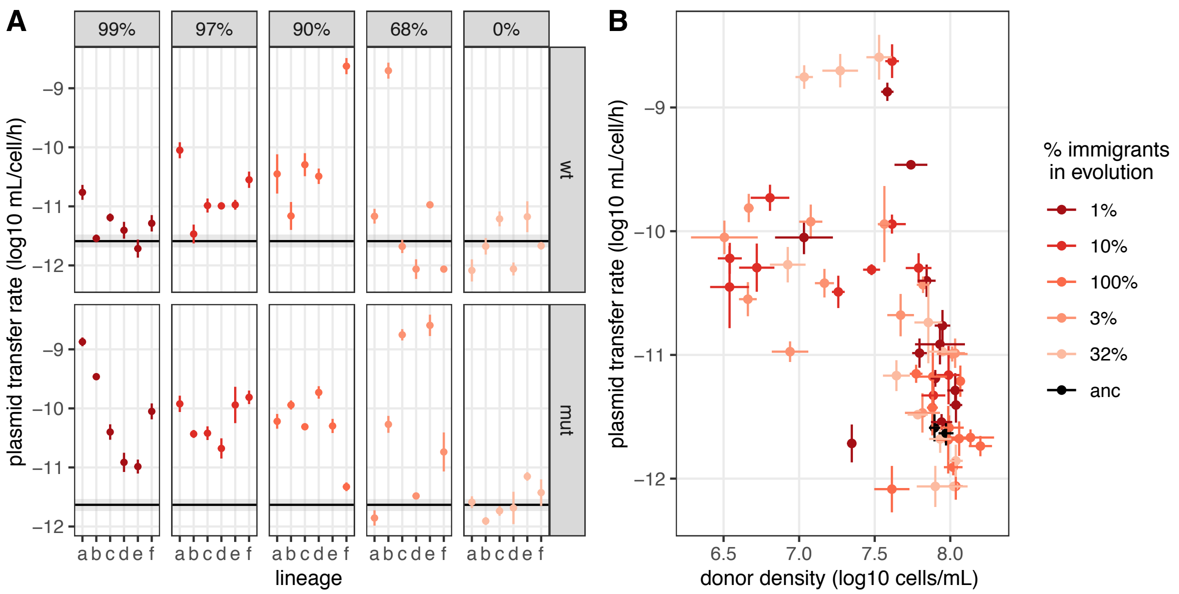

**Figure S2: Midpoint R1 plasmid specific conjugation rates (A) and associated trade-off with donor density (B).** Plasmid transfer rates were measured from plasmid-bearing ancestral hosts towards the standard recipient strain MG1655 $Nal^R$. Dots are geometric averages and lines indicate geometric standard errors (N=3). In A, the black line and shaded area are respectively the geometric average and standard error of ancestral plasmid transfer rate. We observed a strong effect of immigration treatment on evolved transfer rates (transfer rate ~ immigration, $F_{5,218}$=19.0, $p$=1.06.$10^{-15}$). Specifically, treatments with ≥68% immigration had significantly increased transfer rates compared to the ancestor and no immigration treatments (Tukey test, $p < 4.10^{-5}$ for all) but were not significantly different from each other. In B, transfer rates measured from ancestral hosts are shown as a function of donor host cell density at the end of the conjugation assay. Host fitness correlated significantly with plasmid transfer rates (estimate=-0.32±0.03, $r^2$=0.34, $p$<$2.10^{-16}$)

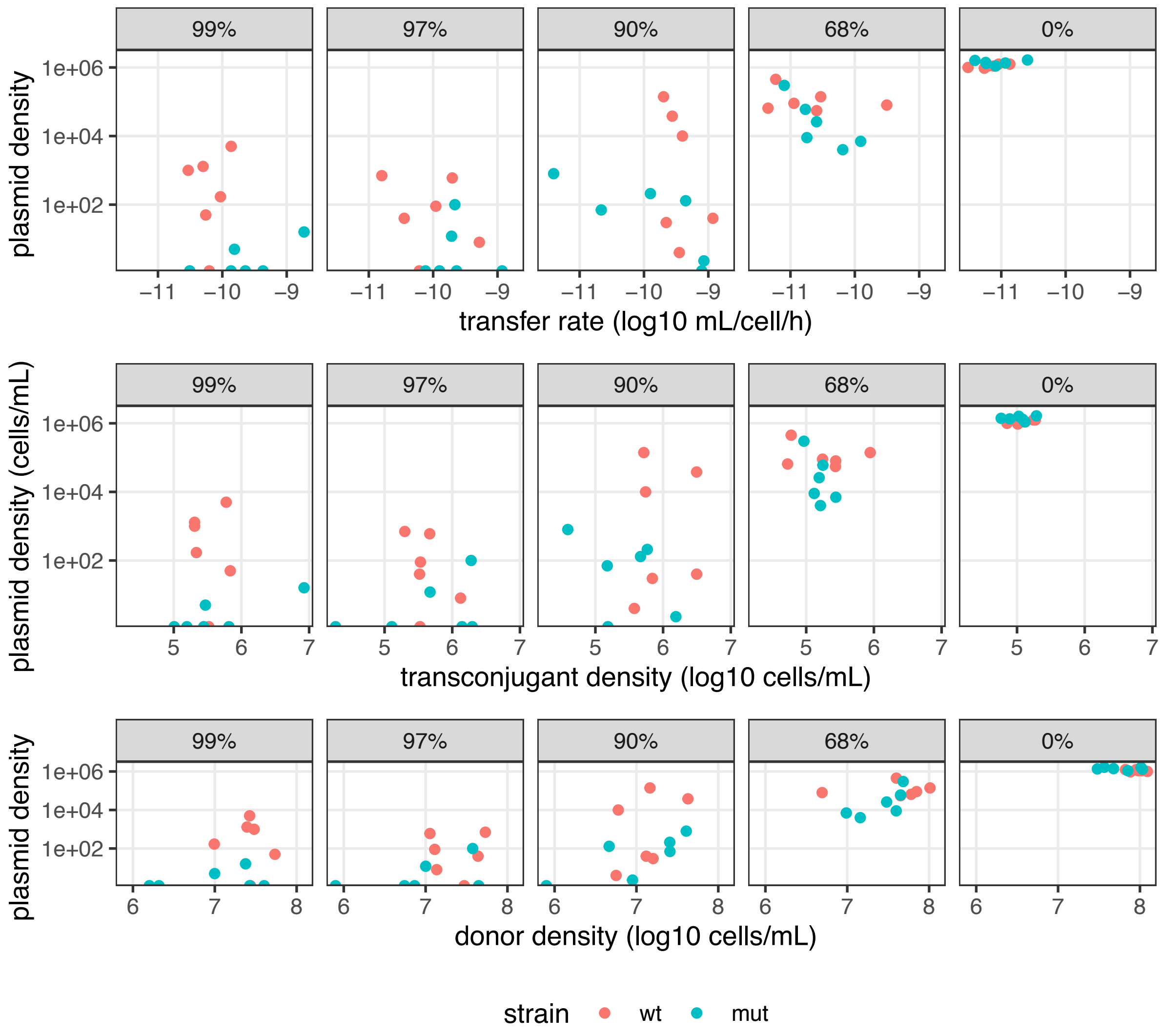

**Figure S3: Relationship between evolved clone phenotypes and plasmid maintenance in evolving populations.** Plasmid-bearing cell population density in evolving populations at day 19 is plotted as a function of the phenotypes of corresponding isolated clones from day 19: transfer rate (top), transconjugant density (middle) and donor density (bottom). Evolved transfer rate correlates negatively with the density of plasmid-carrying cells in the evolution experiment (plasmid density ~ treatment + $\log_{10}$ transfer rate, transfer rate effect = -0.70±0.35, $F_{1,55}$=4.1, $p$=0.048); and donor host density in conjugation assays correlates positively with plasmid maintenance during the evolution experiment (plasmid density ~ treatment + $\log_{10}$ donor density, transfer rate effect = 1.43±0.39, $F_{1,55}$=13.3, $p$=0.0006), suggesting vertical transmission was more important than horizontal transmission for plasmid maintenance.

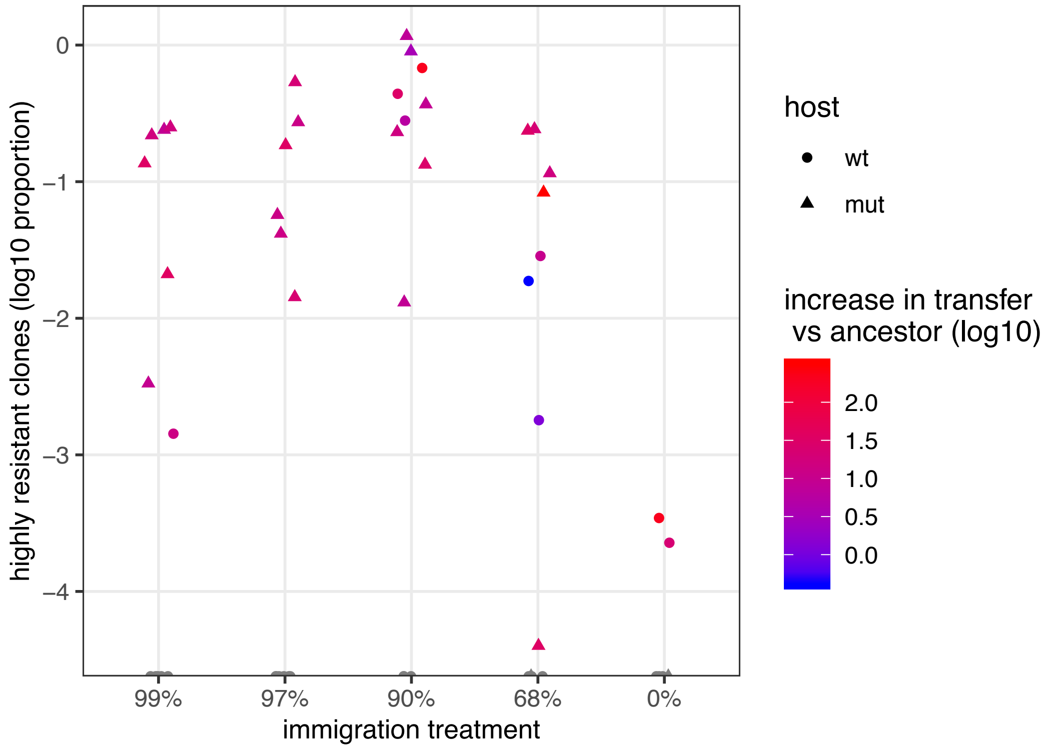


**Figure S4: Proportion of highly resistant plasmids after selection for transfer.** The proportion of plasmid-bearing cells able to grow in the presence of 0.5g/L Amp is shown as a function of immigration treatment in 9 day evolved populations. For each population, a clone was picked randomly and its transfer rate was measured (N=2). Dot color indicates the ration of evolved transfer rate to ancestral R1 transfer rate ($\log_{10}$).

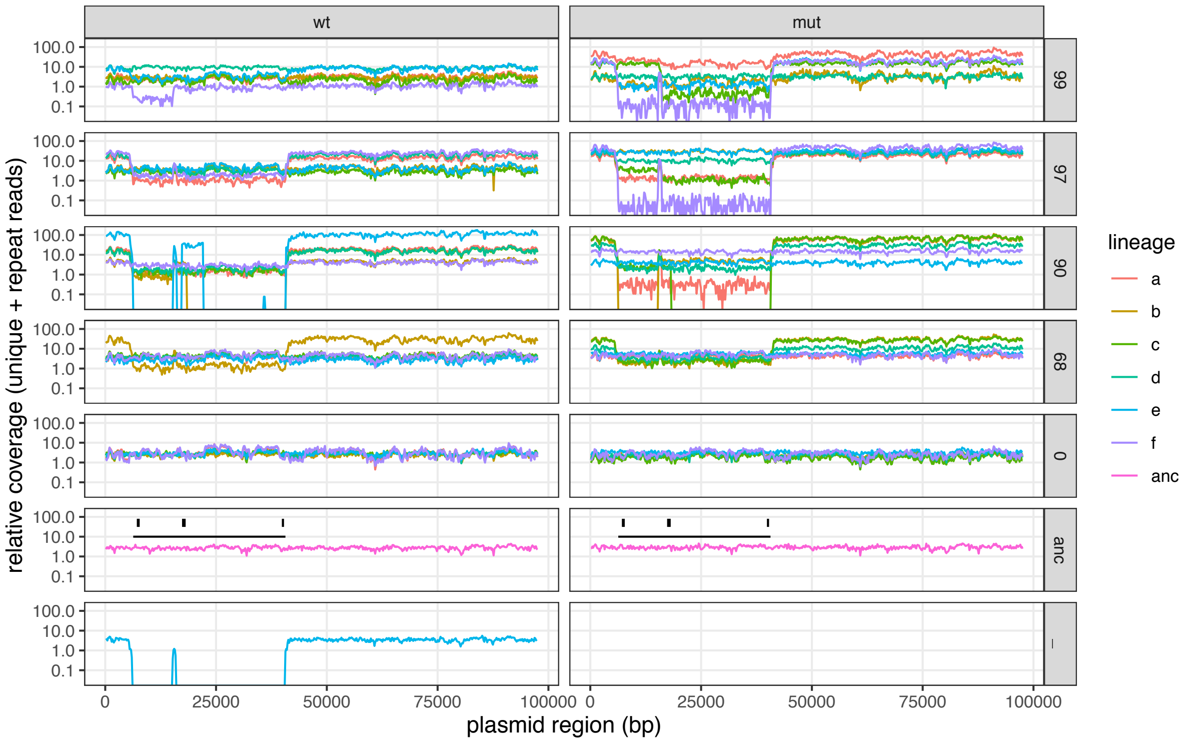


**Figure S5: Variation in sequencing coverage across R1 plasmid map.** Relative coverage of sequencing reads is shown for all clones across $R1_{wt}$ sequence map. Relative coverage was measured as the sum of coverage of both unique and repeat reads, divided by the overall average coverage of reads mapped to the chromosome. Black rectangles above the ancestor graph indicate the position of genes conferring antibiotic resistance (from left to right: kanamycin, ampicillin and chloramphenicol), and the black line indicates the whole resistance determinant region (bordered by insertion sequences). In the plasmid-free control treatment (bottom), reads mapping to the plasmid were detected for lineage w_e, indicating contamination by a plasmid missing the whole resistance determinant. All other clones retained at least the *bla* gene conferring Amp resistance, as they were obtained by plating on Amp-containing medium.

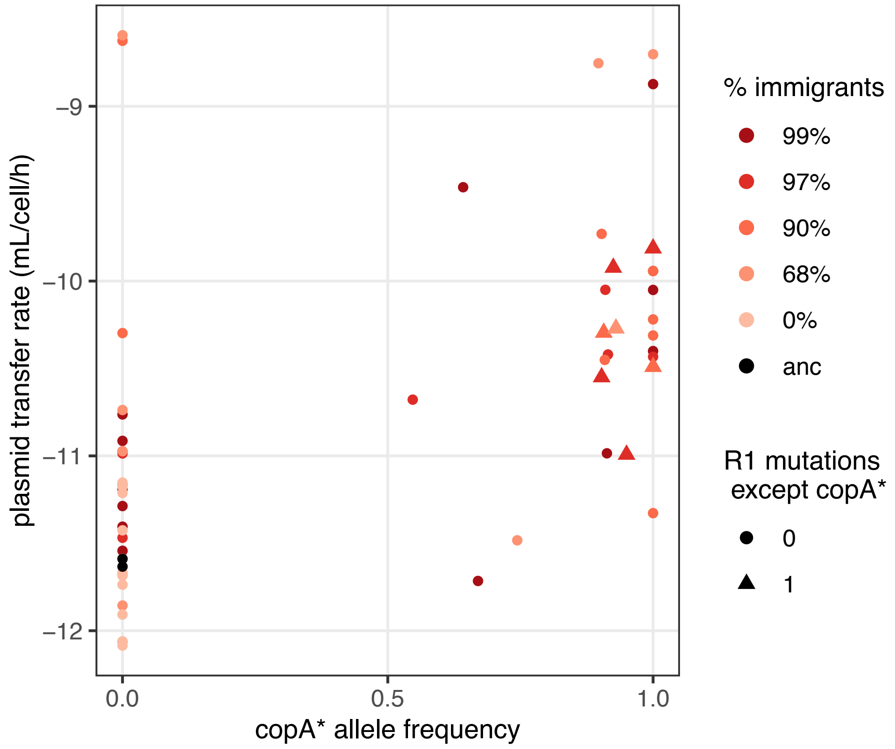


**Figure S6: Effect of *copA** variants on evolved plasmid transfer rate from the ancestral host.** Evolved rate is shown as a function of *copA** mutation frequency in the evolved clone. Triangles indicate clones for which no other mutation than *copA** was present on the plasmid.

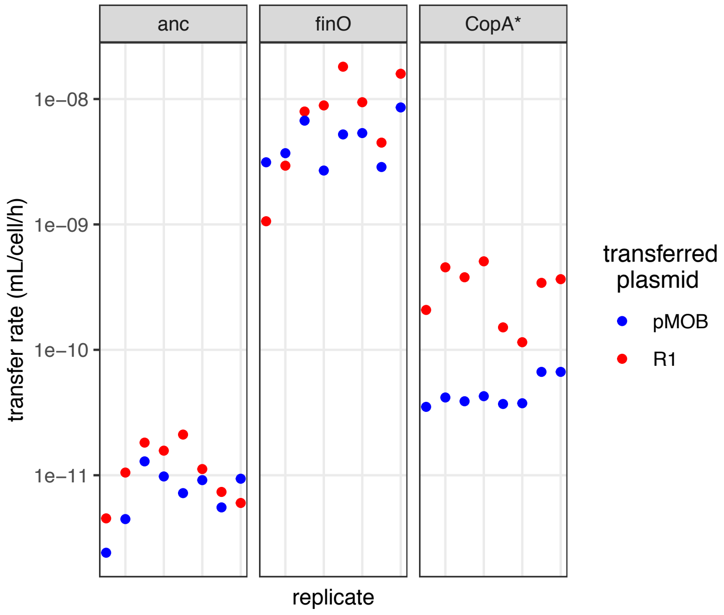


**Figure S7: Impact of evolved *copA** variants on plasmid mobilisation *in trans*.** R1 transfer was measured together with mobilisation of pMOB carrying R1 *oriT*. Increased transfer operon expression will increase transfer of both plasmids, whereas increased *oriT* copy number will only increase R1 transfer.

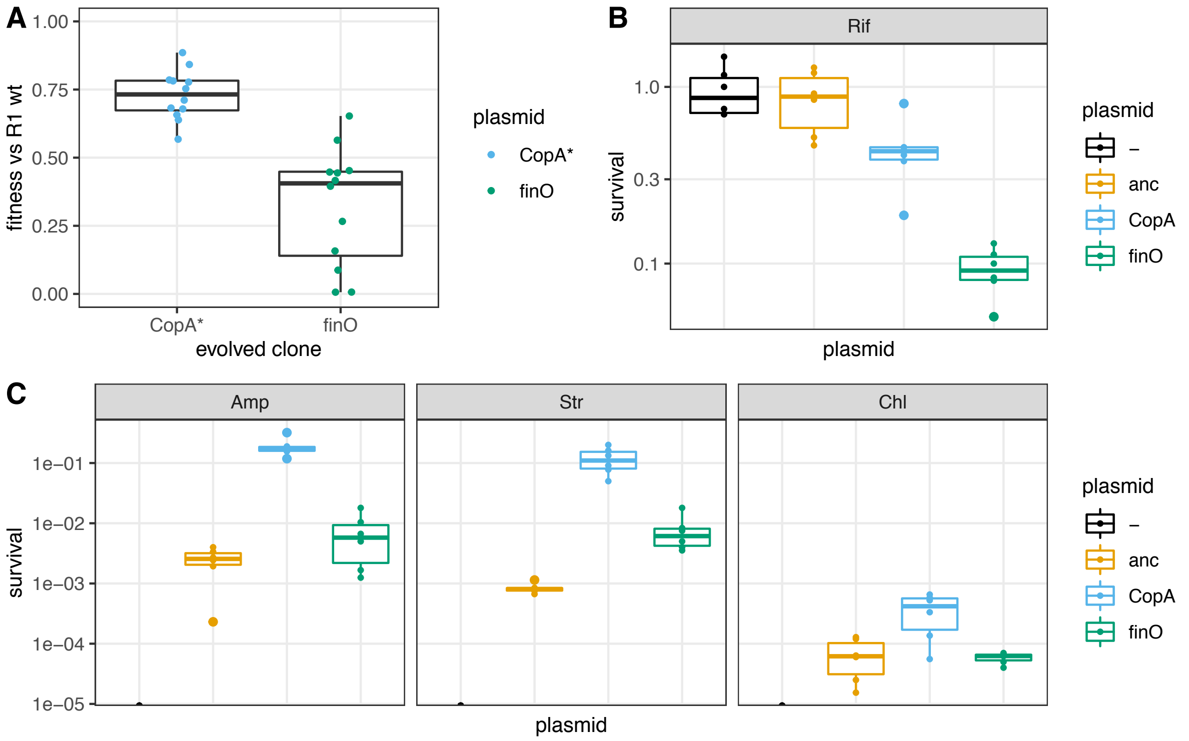


**Figure S8: Fitness and survival of cells carrying R1 variants.** We characterised two variants each with a single mutation (see Table S2): one *copA** variant (plasmid from evolved clone w90d) and one *finO* variant (that has derepressed transfer without copy number mutations, plasmid from evolved clone m68e_t12), in a standard host. A: Relative fitness in competition with $R1_{wt}$ -bearing clones, in the absence of antibiotics (*N=12*). B: Survival in the presence of 2 mg/L rifampicin. Cells carrying each variant survived less compared to $R1_{wt}$ carrying cells, with $R1_{finO}$ conferring the strongest cost. C: Survival in the presence of high doses of antibiotics to which R1 carries resistance determinants. Antibiotics tested were ampicillin 500 mg/L (Amp), streptomycin 200 mg/L (Str) and chloramphenicol 600 mg/L (Chl). In B and C, dots indicate individual populations; the centre value of the boxplots is the median and boxes denote the interquartile range (*N=8*).

**<u>Table S1</u>: Detail of characterized R1 clones.** Midpoint clones were isolated between passage 16 and passage 19: one Amp resistant clone per lineage was chosen randomly for each lineage. When possible, clones from passage 19 were isolated but in lineages where plasmid-bearing cells were already extinct at passage 19, one clone was isolated from populations from passages 17 or 16. Two additional clones from the m68e lineage were isolated from different timepoints and sequenced; the m68e_t12 clone was found to carry only one mutation in *finO* (see Table S2). W90d clone carried only one mutation in *copA*. Both clones were used for further assays (Figure S7 and Figure S8).

**<u>Table S2</u>: Sequence variants detected in evolved clones.** A table was generated with breseq option gdtools COMPARE, then manually cleaned. For the plasmid, two types of mutations were discarded. One was a point mutation 58157 C>T, present in the ancestor and all evolved plasmids. The second was a cluster of complex mutations in positions 87616-87664, within *traD* CDS. The ancestor and many but not all evolved clones carried various 9nt insertions or deletions when compared to KY749247. A similar 9bp insertion is also found in AY684127 accession of *traD* sequence. For the chromosome, variants with less than 100% allele frequency were discarded.
